# A novel supplemental circadian feedback loop in CA1 mediates mood-related behaviors

**DOI:** 10.1101/2023.01.13.524012

**Authors:** Xin-Ling Wang

## Abstract

Traditional circadian clock feedback loops include positive branches and negative branches. *Per* genes belong to the negative branches. There are three subtypes of *Per* genes named *Per1, Per2* and *Per3*. The relationship among these subtypes has been rarely reported. In this study, we aimed to explore the action between *Per1* and *Per2* genes, which are known to be critical in the pathogenesis of mood disorder. We revealed that *Per1* has a positive action on the expression of *Per2*, while *Per2* shows a negative effect on *Per1* expression. This forms a novel feedback loop. Besides, both knockdown and over-expression of *Per1* exhibit a pro-depressive effect, indicating a potential mediation in the pathogenesis of major depressive disorder. Correspondingly, knockdown of *Per2* induces mania-like behavior, while, over-expression of *Per2* produces a pro-depressive effect, suggesting its involvement in the pathophysiology of bipolar disorder. This research may provide an advance in the differential diagnosis between the two diseases in the future.

**Highlights:**

1. *Per1* promotes the expression of *Per2*, while *Per2* inhibit the expression of *Per1* in CA1, forming a negative feedback loop.
2. Both knockdown and over-expression of *Per1* in CA1 induce depression-like behaviors, while *Per2* involves in both mania and depression-like behaviors.

## Introduction

Traditional transcription feedback loop of molecular circadian system includes positive arms and negative arms. CLOCK and BMAL1 complex actives the transcription of *Per* and *Cry*, meanwhile, the PER and CRY forms complex and inhibits the transcription of *clock* and *bmal1* genes, forming the core clock negative feedback loops. Besides, there are three subtypes of *Per* genes, including *Per1, Per2* and *Per3*. Until now, there are rare reports on the relationship between *Per1* and *Per2* [1], and they both participate in mood disorders [2, 3]. It was revealed that mPER1 and mPER2 interact with each other [4, 5]. Zheng *et al*. deduced that they may have complementary effect in the function of maintaining circadian clock [6]. Otherwise, Brenna *et al*. reported that PER2 with CREB forms as a complex that may promote the transcription of *Per1* [7]. Until now, there is not a consistent conclusion on the interaction between *Per1* and *Per2*.

In our former study, we reported that *Per1* knockdown in CA1 induced depression-like behavior in rats [3]. Furthermore, in our recent study, we have found that *Per2* knockdown in CA1 promotes mania-like behavior, in contrast, *Per2* over-expression produces depression-like behavior [8]. However, the action between *Per1* and *Per2* and the roles of their production in MDD or BD have not been revealed. In this study, we reported a novel feedback loop of *Per1* and *Per2* and their different roles in mood-related behaviors, which indicates the roles of them in the pathogenesis of major depressive disorder and bipolar disorder.

## Materials and Methods

### Animals

Male Sprague Dawley (SD) rats (240-260 g) were purchased from the Ji’nan Pengyue Laboratory Animal Breeding Company and housed in groups of three under a reverse 12 h/12 h light/dark cycle (8:00 light on (zeitgeber time 0) and 20:00 light off (zeitgeber time 12)) with free access to food and water. The room temperature was maintained 23±1°C and humidity was 50±5% controlled.

All of the animal procedures were performed in accordance with the National Institutes of Health Guide for the Care and Use of Laboratory Animals and were approved by the Ethics Committee of Shandong University School of Basic Medicine (Ethical approval number: ECSBMSSDU2022-2-51).

### Drugs

#### Intracranial microinjections

After the rats were anaesthetized by isoflurane gas, we microinjected Lentivirus (LV)-Per2/LV-Per1 (1ul per side) or LV-Control (1ul per side) bilaterally into CA1 (anterior/posterior, −4.3 mm; medial/lateral, ±2.0 mm; and dorsal/ventral, −2.0 mm with Hamilton syringes connected to 30-gauge injectors. We infused within 5 minutes and kept the injector in place for additional 3 minutes to allow diffusion.

### Sucrose preference test (SPT), forced swim test (FST), Open field test (OFT) and elevated plus maze (EPM)

The procedures were based on our previous studies [3, 8].

### Western Blot

Rats were killed at zeitgeber time (ZT)2-ZT4, CA1 brain tissues were collected and proteins were extracted for western blot. The western blot procedure was referred to our previous studies [3]. GAPDH was used as the internal reference.

### Quantitative real – time PCR (QRT-PCR)

Rats were killed at zeitgeber time (ZT)2-ZT4, CA1 brain tissues were collected and RNAs were extracted for qRT-PCR. We conducted qRT-PCR referred to our former study [9]. The primers designed for *rPer1* and *rPer2* were shown as follows: *rPer1* forward primer, 5*’* - CCT CGATGT AAC GGC TTG TGT - 3 *‘*; *rPer1* reverse primer, 5*’* - GGAAGA GCT CCC ACC TTG TTC - 3 ‘; *rPer2* forward primer, 5*’* - GGAAGT TCT GGC TGC ACA TAC - 3 *‘*; *rPer2* reverse primer, 5*’* - AGA CCC TGT GCT CTC AGA AGA - 3 *‘*;

### Statistical analysis

The data were expressed as mean ± SEM. Statistical analyses were performed using Prism 5 software (GraphPad). The statistical analyses of data were done using unpaired two-tailed Student’s *t*-test, which can be referred to the figure legends. Values of *p* < 0.05 were considered statistically significant.

### Results

#### 1. Over-expression of *Per2* inhibits Per1 expression levels and shows pro-depressive-like effect in rats

In order to explore the action of *Per2* on *Per1*, we over-expressed *Per2* levels in CA1 by infusing LV-*Per2* into CA1 region of rats and decapitated the rats 12 days after the surgery (Fig. 1a). We showed the infusion site and the transfection cells in CA1 through immunofluorescence (Fig. 1b). Western blot analysis showed that Per2 levels were significantly increased in the LV-*Per2* group, compared with the control group (Fig. 1c-d, *p* < 0.05). On contrast, we found that Per1 levels were significantly decreased in the LV-*Per2* group, compared with the control group (Fig. 1c, e, *p* < 0.05). This result demonstrated that over-expressed Per2 may have an inhibited effect on *Per1* expression. What’s more, over-expression of *Per2* in CA1 showed a depression-like effect in rats, which can be referred to our former paper [8].

**Figure 1.**
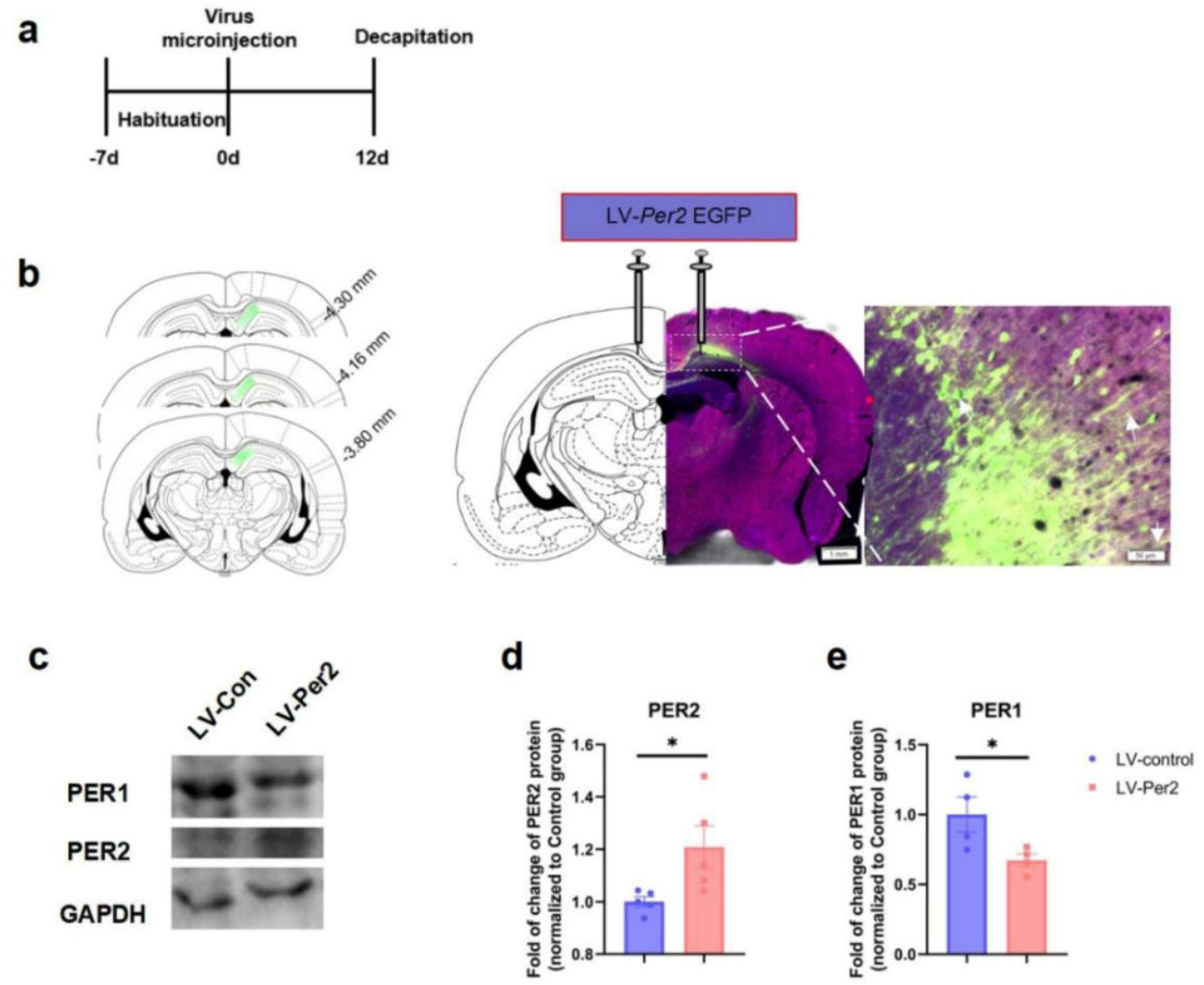
Over-expression of *Per2* in CA1 inhibit the expression of PER1. a. Experimental timeline. b. Immunofluorescence showed the infusion site of LV in hippocampal CA1 region. c. Westernblot analysis of over-expression of Per2 in CA1. Unpaired *t* test, \**p* < 0.05, compared with the LV-control group.

#### 2. Over-expression of *Per1* in CA1 promotes the expression of *Per2* and exhibits a pro-depressive-like action

In order to reveal the effect of *Per1* on *Per2*, we micro-injected LV-*Per1* into CA1 region in rats. 12 days later we killed the rats, collected the CA1 tissue and extracted the RNAs (Fig. 2a). The infusion site and the transfection cells in CA1 through immunofluorescence (Fig. 2b). We performed qRT-PCR and showed that *Per1* levels were significantly elevated in the LV-*Per1* group, compared with the control group (Fig. 2c, *p* < 0.001). What’s more, we found that *Per2* levels were also significantly up-regulated in LV-*Per1* group, demonstrating a promoted effect of *Per1* on *Per2* expressions (Fig. 2d, *p* < 0.001).

**Figure 2.**
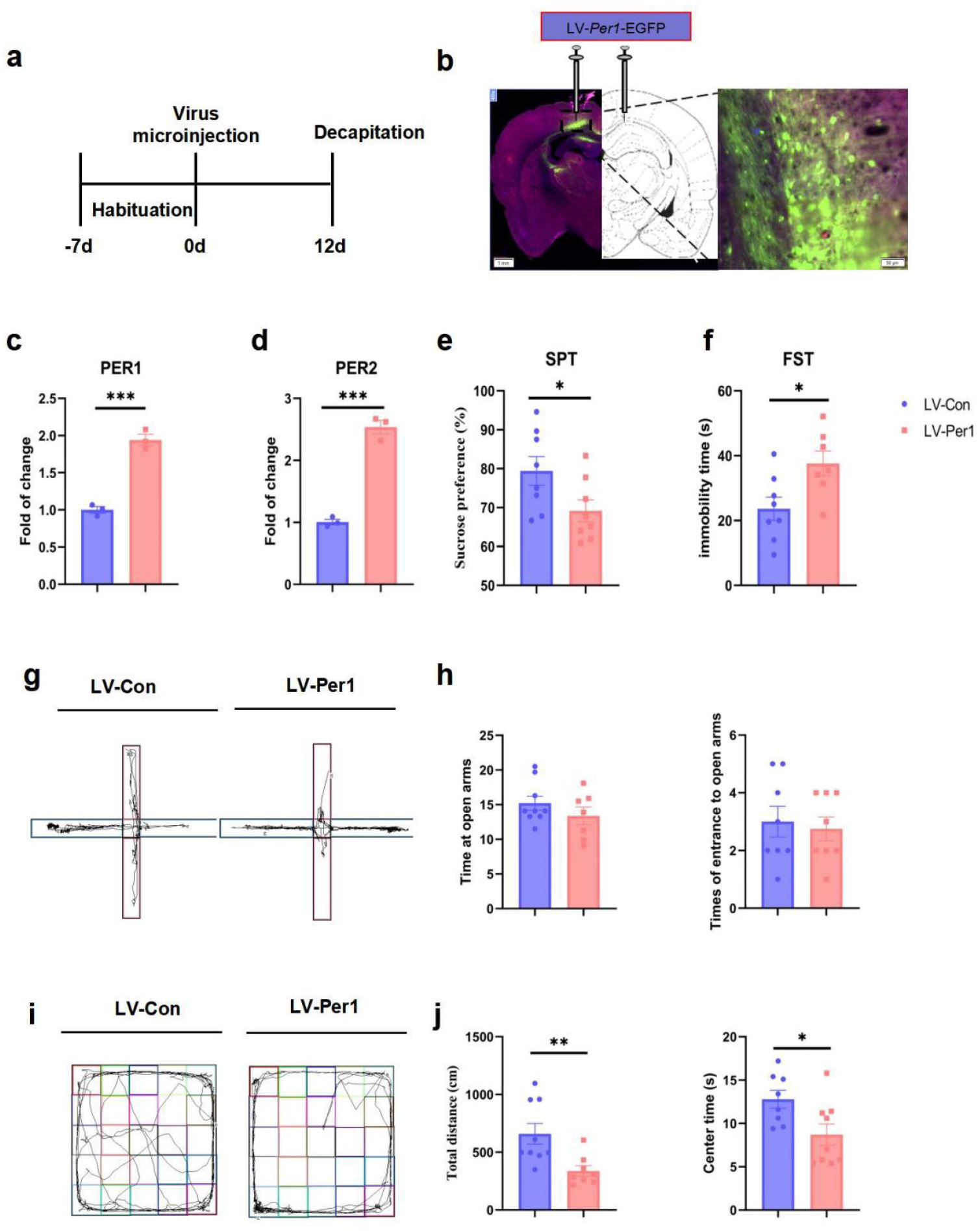
Over-expression of *Per1* in CA1 promotes the expression of *Per2* and depression-like behavior. a. Experimental timeline. b. Immunofluorescence showed the infusion site of LV in hippocampal CA1 region. c. qRT-PCR showed the influence of *LV-Per1* on the expression of Per1. d. qRT-PCR showed the influence of LV*-Per1* on the expression of *Per2*. e. Sucrose preference test. f. Forced swimming test. g. The representative figure of elevated plus maze test. h. The left panel showed time in open arms of both groups. The right panel showed times of entrances to open arms. i. The representative figure of open field test. j. The left panel showed total distances of both groups. The right panel showed time spent in center zones. Unpaired *t* test, \**p* < 0.05, \*\**p* < 0.01, \*\*\**p* < 0.001, compared with the LV-control group.

We also performed a paralleled study. 12 days after the micro-injection surgery, we conducted sucrose preference test (SPT), forced swimming test (FST), elevated plus maze test (EPM) and open field test (OPT) during the consecutive days from Day 12 to Day 15. Sucrose preference test showed that the sucrose preference values were significantly down-regulated in the LV-*Per1* group, compared with the control group (Fig. 2e). Besides, forced swimming test showed that immobility time was significantly prolonged in the LV-*Per1* group (Fig. 2f), showing depression-like behaviors. In the elevated pulse maze test, we didn’t find any significant difference between the LV-*Per1* group and the control group (Fig. 2g-h). In addition, the open field test revealed that the total distances were significantly decreased in the LV-*Per1* group (Fig. 2 j, left panel, *p* < 0.01), what’s more, time spent in center zones was also significantly reduced in this group (Fig. 2 j, right panel, *p* < 0.05), compared with the control group, showing anxiety-like behaviors.

## Discussion

In this study, we demonstrated that *Per1* promotes *Per2* expression in transcriptional levels, while Per2 suppress *Per1* expression in protein levels, forming a novel feedback loop of molecular circadian system in CA1 (Fig. 3); Also, we found that over-expression of *Per1* in CA1 produced depression-like behaviors. Combined with our former researches [3], we conclude that both knockdown and over-expression of *Per1* could cause depression-like behaviors in rats. In contrast, knockdown of *Per2* leads to mania-like behavior, while over-expression of *Per2* produces depression-like effect [8]. All these results indicated that *Per1* may mediate the pathogenesis of major depressive disorder and *Per2* may involve in that of bipolar disorder (Fig. 3).

**Figure 3.**
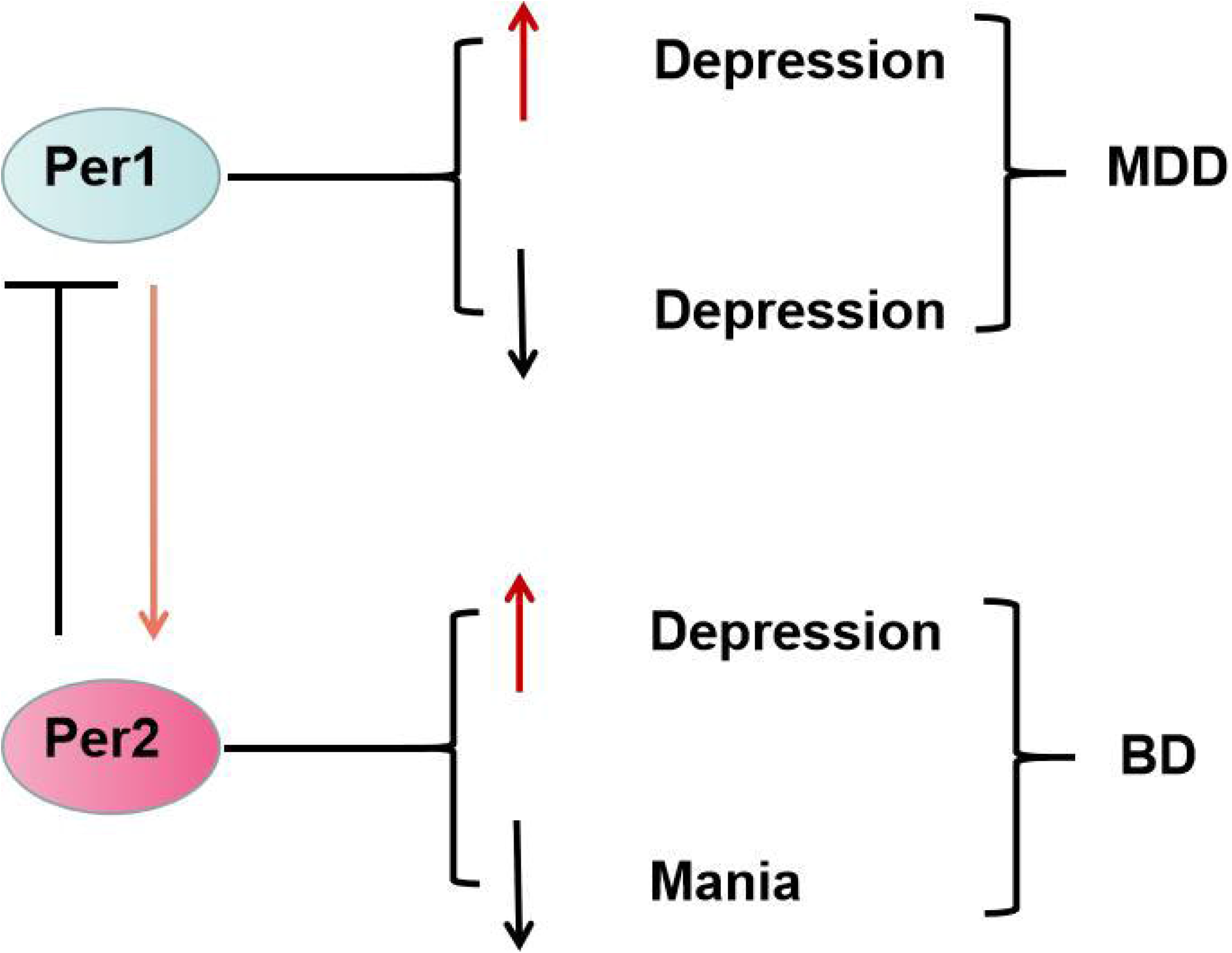
Summary schematic. *Per1* promotes the expression of *Per2*, by contrast, Per2 inhibits the expression of *Per1;* Both knockdown and over-expression of *Per1* in CA1 region leads to depression-like behaviors, which may mediate major depressive disorder. Nevertheless, *Per2* knockdown in this region produces mania-like behavior, meanwhile, *Per2* over-expression in this region promotes depression-like behavior, both of which may mediate the pathogenesis of bipolar disorder.

There have been rare reports on the interaction between *Per1* and *Per2* and their productions. Zheng *et al*. suggested that absence of mPER1 leads to an elevated level of mPER2, indicating that mPER1 may inhibit mPER2 levels in mice SCN [6]. What’s more, Brenna *et al*. reported that PER2 may promote *Per1* expression by combining with CREB complex in SCN [7]. However, our results were contrary to these results, probably due to the different brain regions we researched. We deduced that there might be a tissue-specific effect on the interaction between *Per1* and *Per2*, as our studies are conducted in rat CA1 brain region.

About the phenotype, we demonstrate that *Per1* involves in depressive-like phenotype, which means that it may mediate the pathogenesis of MDD. Meanwhile, *Per2* participates in both mania and depressive-like phenotype, meaning that it may mediate the pathogenesis of BD, in agreement with previous studies [10]. To our knowledge, we are the first to report the relationship of pathogenesis between MDD and BD. In the future, we will make further efforts to explore detailed interacted mechanism of this feedback loop and create novel pharmaceutical aiming at this process for the treatment of BD and MDD.

## Author Contribution

This research was conceived and done by Xin-Ling Wang herself. Furthermore, this manuscript was composed by Xin-Ling Wang.

## Acknowledgement

This study was supported by the National Natural Science Foundation of China (NSFC82201682), Natural Science Foundation of Shandong Province (ZR2021QH282) and the Fundamental Research Funds of Shandong University (2020GN095) to Xin-Ling Wang.

## Conflict of Interests

The author declares that there are no conflict of interests.

## Notes

### Competing Interest Statement

The authors have declared no competing interest.

